# Gaze-centered spatial coding of touch on a hand-held tool

**DOI:** 10.1101/2025.04.04.647167

**Authors:** Lefteris Zografos Themelis, W. Pieter Medendorp, Luke E. Miller

**Affiliations:** Laboratory of Cognitive Neuroscience and Sensorimotor Control, University Mental Health Neurosciences and Precision Medicine Research Institute "COSTAS STEFANIS", Athens, Greece; Donders Institute for Cognition and Behavior, Radboud University, Nijmegen, 6525GD,The Netherlands

**Keywords:** Tool use, sensory projection, embodiment, sensorimotor, somatosensory

## Abstract

Humans possess the remarkable ability to project tactile sensations outside their body and onto a hand-held tool that they are using. Despite nearly a century of research, the computations underlying this projection have not been adequately addressed. In the present study, we employed model-driven psychophysics to fill this gap. We hypothesized that tool-based sensory projection involves the remapping of touch from sensory feedback in the hand into an egocentric coordinate system. We first formalized the computational steps underlying tactile remapping. We designed a novel tool-sensing experiment that allowed us to rigorously test this model. In this task, participants contacted an object with a hand-held rod and then judged whether the object was above or below where they were currently looking. This comparison would only be possible if touch on the tool was projected outside the hand. Crucially, both object location and gaze position varied independently, allowing us to characterize the hand-to-space-to-gaze remapping process. Model-based curve fitting provided strong evidence that all participants in our task projected touch outside their body and into gaze-centered coordinates. Crucially, the resolution of this projection was similar to what has been found for touch on the body. These findings provide the first step towards characterizing the computations underlying the spatial projection of touch on external objects, highlighting the incredible versatility of the sensorimotor system.

## Introduction

In 1911, the neurologists Head and Holmes proposed that humans can project their sense of locality outside their body and onto hand-held implements (1). Research since then has indeed found a strong sensorimotor link between the body and tools (2-10). For example, humans can use tools to accurately sense objects in the environment (11), including their surface texture (12-14) and distance (15). Furthermore, they can as accurately localize touch on a tool as they would on their own body (16-21), perhaps by repurposing body-based neural processing for tactile localization (22-26). Little is known, however, about the computations by which the sensorimotor system projects the sensation of touch on an object outside the body (1). We aimed to address this in the present study.

Let us start by asking a seemingly simple question: How might someone direct their gaze towards the location of an object touched by a tool? To gain computational traction on this question, we can first address how this would happen when touch is on the hand. Looking towards an object touching the hand requires knowing the touch’s precise location relative to gaze, i.e., in external space. Since cutaneous signals are skin-based, they must first be integrated with proprioceptive information about the hand’s location in external space (27-29). Computationally, this amounts to remapping touch from a skin-based signal into a coordinate system anchored to the hand (hand-centered tactile remapping) (30). Looking towards this location requires further re-mapping (31, 32), where this hand-centered representation is transformed relative to the current location of gaze (gaze-centered tactile remapping).

We propose—and experimentally verify the idea—that tactile remapping is a candidate computation for projecting touch outside the body. Returning to our guiding example: Since a tool does not have intrinsic sensors, information about touch location must first be derived indirectly from mechanical signals (16), such as vibrations or torque. Proprioceptive information about tool posture (33-35) is therefore necessary to remap these mechanical signals outside the body to the tool’s location in external space (perhaps anchored to the hand). Looking towards this location would require further remapping the projected touch to the location of gaze (gaze-centered remapping).

In the present study we characterize these steps, providing a computational basis for projecting touch outside the body. We developed a 2AFC task that required participants to remap tactile sensations during tool-sensing from hand-centered to gaze-centered coordinates (Figure 1A). In the task, participants used a carbon fiber rod to hit an object that was placed above their hand and judge whether the object was located above or below their current gaze position. We then used model-driven curve fitting to analyze participant’s responses. To foreshadow our results, we indeed found that tool-users accurately remap projected touch into gaze-centered coordinates, and do so with a similar precision as touch on the body.

**Figure 1.**
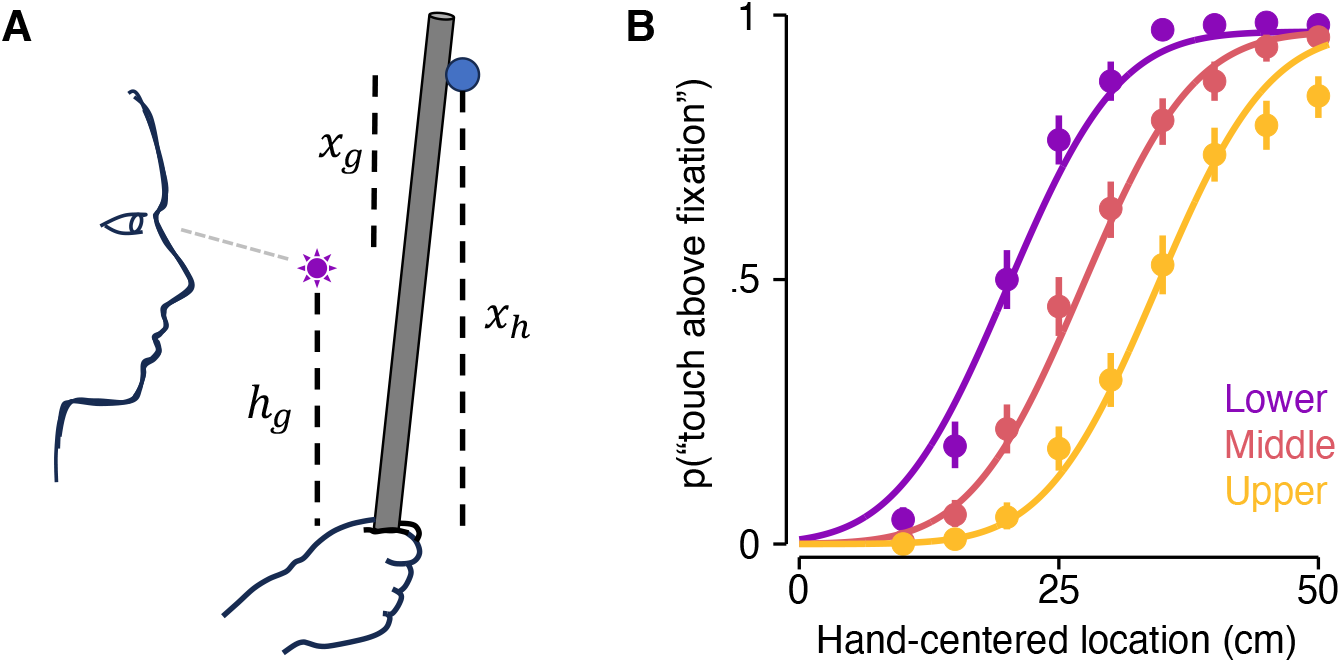
Remapping paradigm and group-level results. (A) Localizing the location of the object (blue circle) relative to the position of gaze (purple sun) requires remapping touch into gaze-centered coordinates. (B) Model fit for the group-level responses for each of the three gaze positions (hand-centered coordinates). Errors bars reflect the bootstrapped 95% confidence intervals.

## Materials and Methods

### Participants

Twelve right-handed participants (seven females) averaging 24.3±2.1 years of age participated in this study. All participants had normal or corrected-to-normal vision and no history of neurological disorders (e.g., psychosis), chronic pain, or motor disturbances. All signed an informed consent before the start of the experiment. The experiment was approved by the local ethics committee of the Faculty of Social Sciences at Radboud University.

### Experimental Setup

Participants sat comfortably in a chair, holding a 73 cm carbon-fiber rod in their right hand behind a semi-translucent vertical occlusion board. Their forearm was placed on an armrest, which minimized fatigue and excessive arm movements during the experiment. Their head was placed on a chinrest attached to the uppermost portion of the occluding board. The height of the chair was adjusted so that the participant’s chin could rest comfortably in the chinrest while they sat upright. A strip of LED lights was placed vertically behind the occluding board to serve as fixation points (25 cm horizontal distance from the eyes). These LEDs could shine at three different colours (red, blue, green), and were clearly visible to participants when turned on.

The experimental task (see below) involved hitting an object (16 cm horizontally-oriented carbon-fiber tube) with the aforementioned rod. A custom-built motorized system was used to control the placement of this object in the vertical plane (functional range: 0–60 cm) with millimeter precision. This system remained behind the occluding board, thus preventing participants from ever seeing the object’s current spatial location. To ensure that participants did not hear the rod-object contact, participants wore noise-cancelling headphones that played white noise. Participants used a button box placed below their left hand to report judgments during the task.

### Touch localization paradigm

The participant’s task was to determine whether the location of the object struck with the rod was above or below their current gaze location (i.e., fixation point). On a given trial, the object was placed at one of nine locations (10–50 cm above the hand, in steps of 5 cm) and the fixation point was at one of three locations (25, 30, or 35 cm above the hand). There were thus 27 possible eye-object pairs. The combinations of eye and object locations (in hand-centered coordinates) meant that there was a total of eleven possible gaze-centered object locations (−25–25 cm, in steps of 5 cm) in the experiment.

A typical trial was as follows: The participants began by holding the rod vertically, cocked slightly back to rest against the side of the occluding board. A red fixation point appeared at one of the three locations, indicating where they were to look. After 750–1250 ms, the fixation turned green to indicate that the participant should hit the object, which was placed at one of the nine locations. After a short delay period (1000– 2000 ms), the fixation turned blue to indicate the participant to report their judgment. Once the judgment was made the fixation disappeared for 500 ms, before starting the next trial.

The total experiment had six blocks of 81 trials each (3 trials per eye-object pair), separated by short breaks. There were a total of 18 trials per eye-object pair, with 486 total trials in the experiment. The experiment lasted ~1.5 hours in total.

### Model-driven curve fitting

If the tool-user projects their tactile sensations into gaze-centered space (Figure 1A), their perception should vary lawfully as a function of gaze position in the following way:

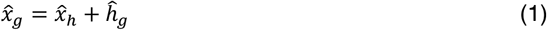

Where 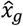 is the estimated location of touch in gaze-centred space, 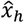 is the projected hand-centred location of touch, and 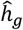 is the estimated location of the hand relative to gaze. Following this equation, we used model-driven curve fitting to determine whether tool-users indeed projected touch on the rod outside the body and into gaze-centered coordinates.

We fit each participant’s judgments in the three gaze-centered conditions with psychometric curves (Cumulative Normal). These functions modelled the probability that the participant reported the object was above the fixation, given the specific hand-centered object location,

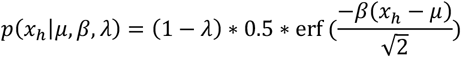

where *x*_*h*_ is the actual stimulus location in hand-centered coordinates, *erf* is the complementary error function, *λ* is the lapse rate, *β* is the slope, and *μ* is the threshold of the curve. Note that, in our experiment, the unit of gaze position was always measured relative to the hand. We performed curve-fitting for touch location in both hand-centered and gaze-centered coordinates. However, because each produced identical estimated parameters, we mainly focus our manuscript on the curve-fitting in hand-centered coordinates. The just noticeable difference of the curve corresponds to the difference between where the curve crosses 0.75 and 0.5.

The threshold was the only parameter in our curve fitting we hypothesized to vary as a function of eye position. It was therefore the focus of our modelling (see below). The two models fit in our study are as follows:

### Projection model (Model P+)

This model assumes that touch on the rod is remapped into gaze-centered coordinates following Equation 1. If this transformation is free from bias, then the threshold *μ* of the psychometric curve corresponds to the position of gaze *g* (in hand-centered coordinates). However, prior research has uncovered two types of biases that would affect the participant’s pattern of judgments (16, 22): (i) A non-zero gain (α) on the relationship between actual and perceived touch location in hand-centered space, 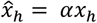; (ii) A misestimate (η) of the hand relative to the center of gaze, 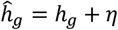.

Taking these two biases into account, the threshold of the psychometric curve corresponds to:

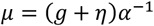

where *α*^−1^ transforms all units into hand-centered coordinates normalized by the gain of the tool-to-space transformation. The model fitting had four free parameters with the following constraints: the slope *β* [0, 1], the distance misestimation *η* [-10, 10], the gain *α* [0.5, 1.5], and the lapse rate *λ* [0, 0.1]. All curve-fitting was performed with the Palamedes Toolbox in Matlab 2022 as well as custom scripts for constrained parameter optimization (using fmincon).

### Non-projection model (Model P-)

In this model, participants do not modulate their judgments based upon gaze position. We assume that participants use a consistent but unknown strategy when making their judgments, and therefore the data across gaze conditions is best fit by a single psychometric curve. The model fitting had three free parameters with the following constraints: the slope *β* [0, 1], the threshold *μ* [5, 45], and the lapse rate λ [0, 0.1].

### Model comparison

We used the Bayesian Information Criteria (BIC) to adjudicate between the above two models for each participant,

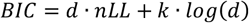

where *d* is the number of datapoints, *nLL* is the negative log-likelihood of the model fit, and *k* is the number of free parameters. For each model, a lower BIC corresponds to a better fitting model, and the difference between them (Δ*BIC*) can be used to objectively compare fitting performance. A Δ*BIC* greater than six corresponds to a significantly better fit for Model P+ compared to Model P-.

## Results

We first sought to confirm that participants remapped touch on the hand-held rod into gaze-centered coordinates. As hypothesized, the spatial judgments of participants indeed varied as a function of gaze position. Both the group-level data (Figure 1B) and the individual-participant data (Figure 2) was well-fit by Model P+. Indeed, Bayesian model comparison demonstrated that Model P+ vastly outperformed Model P- for all participants (median ΔBIC: 57.1; range: 17.1–91.9). Model P+ still outperformed Model P- for 11 out of 12 participants when we assumed the reference frame transformation (Equation 1) included zero biases. These results demonstrate that participants indeed remapped the touch on a tool into gaze-centered coordinates in the present task.

**Figure 2.**
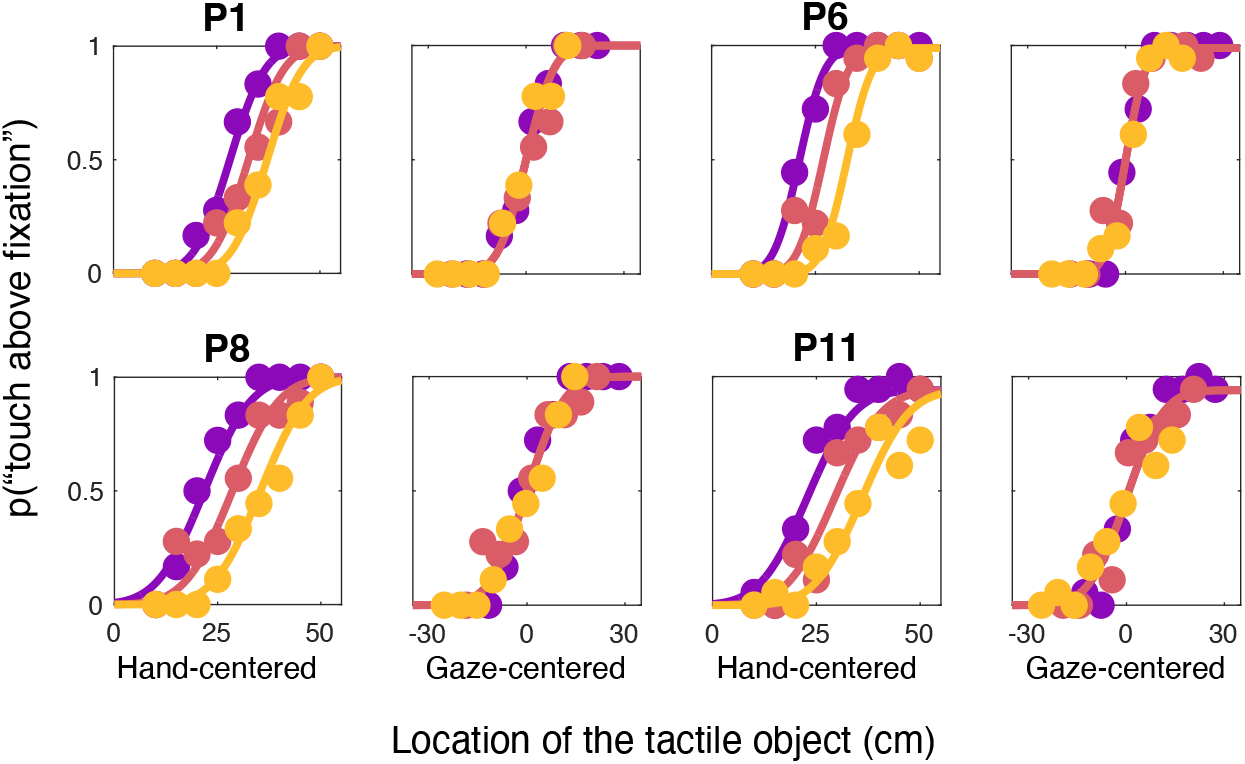
Individual curve fits. Individual curve fits for four randomly selected participants. Fits are presented in both hand-centered coordinates (left panel) and gaze-centered coordinates (right panel). The latter fit collapses into a single curve, demonstrating localization was indeed gaze-centered. As can be seen, the projection model provided a good fit for the spatial judgments of all participants.

Next, we interrogated the model parameters of Model P+ to understand the two spatial biases. The magnitude of the projection gain (*α*) was significantly less than one (mean *α*: 0.88; 95% CI [0.83, 0.93]). The magnitude of this underestimation is within line with what we have seen in prior experiments (11), and is thought to be related to the hand-centered mapping of touch location. Furthermore, we found a significant bias in the estimated hand location (*η*). Specifically, participants perceived the hand and gaze location as closer together than they were (mean *η*: −6.83 cm; 95% CI [−8.51, −5.16]). This is consistent with previous findings of gaze-centered coding of hand position (36).

Finally, we examined the resolution of tactile remapping by quantifying the just noticeable difference (JND) between touch and gaze location. Whereas it is impossible with the current paradigm to tease apart the resolution of each, the JND provides a global measure of the computation’s spatial resolution. The JND was on average near the physical separation between each object location and was highly consistent across participants (mean: 5.52 cm; 95% CI [4.63, 5.42]). Interestingly, this is similar to the JND that has previously been reported for tactile remapping for touch on the arm (37). Overall, this demonstrates that the human sensorimotor system can remap touch on a tool at a fine-grained level.

## Discussion

The present study combined psychophysics and modelling to explore the computations that project touch on a tool into external gaze-centered space. To this end, we developed a novel task that measured participants’ ability to remap tactile sensations during tool-sensing from hand-centered to gaze-centered coordinates (Figure 1A). We found that tool-users can accurately perform this reference frame transformation during tool use. Furthermore, they do with a similar precision as localization on the body.

Tactile remapping is a ubiquitous computation used by the somatosensory system to localize touch on the body (27). Remapping touch into a gaze-centered reference frame appears to be of particular importance. Localization judgments are biased towards the center of gaze (32, 36), even when it is task irrelevant. In accordance with this, parietal cortices readily remaps touch into a gaze-centered reference frame during reaching (38).

Leveraging the importance of gaze-centered coding provided us the opportunity to interrogate the computations underlying perceptual projection. Our paradigm required participants to compare the precise location of touch on a tool with their current position of gaze. Successfully doing so required three computational steps (Equation 1): (i) Remapping from intrinsic (e.g. vibrations on the tool) and proprioceptive information (e.g. posture of the tool in space) to a projected estimate of touch in hand-centered coordinates (16); (ii) computing the hand’s relative position to the center of gaze (36); and (iii) using these two spatial codes to remap touch on the tool to a gaze-centered, external coordinate system.

The neural correlates underlying these transformations are not well understood. A recent model of posterior parietal cortex (PPC) argues that its main function is estimating the states of the body and the environment (39). This estimation unfolds within an anterior-to-posterior (body-to-environment) gradient, particularly in the intra-parietal sulcus. This raises the question of where tool-based reference frames are represented within the body-to-environment gradient.

Several studies have emphasized the importance of PPC for tool sensing and control (5, 25, 40-43). Anterior regions of PPC—near the body-based pole—play a role in proprioceptively estimating the spatial extent of tools (44) as well as localizing touch on a tool (23, 26, 45). Interestingly, localizing touch on a hand-held tool leads to a larger posterior activation spread than localizing touch on the body (26). This may suggest that tools are also represented as somewhat intermediate between the body and environment.

Most previous studies on tool-extended tactile localization have measured it indirectly. For example, using tactile temporal-order judgments (17) or requiring participants to make their localization judgments on a drawing of the rod (16). Only a single experiment that we know of has attempted to measure localization in real space, requiring participants to move a cursor on a screen placed several tens of centimeters away from the tool itself (22). The paradigm we used in the present study goes beyond these limitations, requiring participants to judge the precise relative positions of two points in space, the position of gaze and the external localization of the touched object. This allowed us to characterize, for the first time, the computational process of transformation touch into gaze-centered coordinates.

Tools are often said to be embodied by the sensorimotor system, though such a statement is typically left vague. We have argued that embodiment should be viewed as repurposing a body-based computations—for sensing and/or control—to perform a similar behavior with a tool (11). Indeed, recent work has observed deep similarities between localizing touch on tools and limbs at behavioral (16, 17), neural level (23, 25, 26), and computational (24, 30) levels.

Here we provide another similarity at the computational level: The computation of tactile remapping (27) is repurposed for projecting touch outside the body and onto a hand-held tool (1). Interestingly, the spatial resolution of remapping measured in the present study is consistent with what has been observed for remapping touch on the body (37), lending further support to the idea that tools and limbs share a deep similarity (11). However, it must be noted that this comparison is indirect; future research should directly compare the computational parameters for body- and tool-based sensing.

To conclude, the present study provides further evidence for a computational similarity between limbs and tool. Specifically, we found that tactile remapping—a body-based computation underlying tactile localization—is repurposed for projecting the perception of touch on hand-held tools outside the body. We believe that tool-extended sensing provides a unique opportunity to directly compare the computations used for body-based and tool-based behaviors.

## Author contributions

LZF, WPM, and LEM designed the experiment. LZF and LEM collected the data. LEM performed the modelling. LEM and LZF analyzed the data. LEM and LZF wrote the original draft of the manuscript. WPM provided feedback on the manuscript. LEM supervised the project.

